# The genome sequence of the lesser marbled fritillary, *Brenthis ino*, and evidence for a segregating neo-Z chromosome

**DOI:** 10.1101/2021.12.16.472906

**Authors:** Alexander Mackintosh, Dominik R. Laetsch, Tobias Baril, Robert Foster, Vlad Dincă, Roger Vila, Alexander Hayward, Konrad Lohse

## Abstract

The lesser marbled fritillary, *Brenthis ino* (Rottemburg, 1775), is a species of Palearctic butterfly. Male *B. ino* individuals have been reported to have between 12 and 14 pairs of chromosomes, a much reduced chromosome number than is typical in butterflies. Here we present a chromosome-level genome assembly for *B. ino*, as well as gene and transposable element annotations. The assembly is 411.8 Mb in span with contig and scaffold N50s of 9.6 and 29.5 Mb respectively. We also show evidence that the male individual from which we generated HiC data was heterozygous for a neo-Z chromosome, consistent with inheriting 14 chromosomes from one parent and 13 from the other. This genome assembly will be a valuable resource for studying chromosome evolution in Lepidoptera, as well as for comparative and population genomics more generally.

## 1 Introduction

The lesser marbled fritillary, *Brenthis ino* (Rottemburg, 1775), is a species of butterfly in the family Nymphalidae. It has a Palearctic distribution, is widespread in Europe with variance in local abundance, and can be found as far East as Japan and Siberia. It is monovoltine and feeds on plants in the family Rosaceae, including some species in the genera *Filipendula, Aruncus, Sanguisorba*, and *Rubus*. While most butterflies in the family Nymphalidae, and Lepidoptera more widely, have 31 (or close to 31) pairs of chromosomes (de Vos *et al*. 2020), *B. ino*, along with its sister species *B. daphne* (Denis and Schiffermüller, 1775), has an unusually low chromosome count. Federley (1938) reported male haploid chromosome numbers of 12 and 13 for individuals collected in Finland, consistent with segregating chromosomal fissions or fusions in the population. However, males sampled in Finland and Sweden consistently display 13 chromosome pairs (Saitoh 1987, 1991), and never 12. In Japan, where the subspecies *B. ino mashuensis* (Kono, 1931) and *B. ino tigroides* (Fruhstorfer, 1907) are found, a male chromosome number of 14 has been consistently observed (Maeki and Makino 1953; Saitoh *et al*. 1989).

Currently, there are no genome assemblies for species in the genus *Brenthis* and information about chromosome evolution in the genus is confined to cytological data. Here we present a chromosome-level genome assembly of *B. ino* as well as gene and transposable element (TE) annotations. We also provide evidence for the existence of a segregating neo-Z chromosome that results in variable male karyotypes (13-14 chromosomes) within the Spanish population from which we sampled.

## 2 Materials and methods

### 2.1 Sampling

Three individuals were collected by hand netting in Somiedo, Branña de Mumian, Asturias, Spain (SO_BI_364, SO_BI_375, SO_BI_376) and one in Larche, Alpes-de-Haute-Provence, France (FR_BI_1497, RVcoll12O846) (Table S1). Spanish individuals were flash frozen in a liquid nitrogen dry shipper. The French specimen was dried and, after some days, the body stored in ethanol at -20^*°*^C.

### 2.2 Sequencing

High molecular weight (HMW) DNA was extracted from the thorax of a flash frozen individual (SO_BI_364) using a salting out extraction protocol. In brief, tissue was homogenised in cell lysis buffer using a micro-pestle and then incubated with Proteinase K overnight at 56^*°*^C followed by a further one hour incubation at 37^*°*^C with RNase A before precipitating and discarding proteins. Finally, DNA was precipitated using isopropanol and the resulting pellet was washed with ethanol.

Edinburgh Genomics (EG) generated a SMRTbell sequencing library from the HMW DNA which was sequenced on three SMRT cells on a Sequel I instrument to generate 28.4 Gb of Pacbio continuous long read (CLR) sequence data. From the same HMW DNA extraction, EG also generated a TruSeq library (350 bp insert) and 33.5 Gb of Illumina whole genome (WGS) paired-end reads on a Novaseq 6000. Pacbio and Illumina protocols were followed for library preparation, QC and sequencing.

A second individual (SO_BI_375) was used for chromatin conformation capture (HiC) sequencing. The HiC reaction was done using an Arima-HiC kit, following the manufacturer’s instructions for flash frozen animal tissue. The NEBNext Ultra II library was sequenced on an Illumina MiSeq at EG, generating 4.8 Gb of paired-end reads.

Illumina WGS paired-end reads were also generated for the same individual used for HiC sequencing (SO_BI_375) as well as the French female individual (FR_BI_1497) that did not contribute to the assembly.

Paired-end RNA-seq data (for individual SO_BI_376) was previously generated and analysed in (Ebdon *et al*. 2021) (ENA experiment accession ERX5086186).

### 2.3 Genome assembly

Illumina WGS, RNA-seq, and HiC reads were adapter and quality trimmed with fastp v0.2.1 (Chen *et al*.

2018).

The Pacbio reads were assembled with Nextdenovo v2.4.0 (Hu 2021) using default parameters. Contigs were polished twice by aligning Illumina WGS reads and correcting consensus errors with HAPO-G v1.1 (Aury and Istace 2021). Contigs belonging to non-target organisms were identified using blobtools v1.1.1 (Laetsch and Blaxter 2017) and subsequently removed. Duplicated regions (haplotigs and overlaps) were identified and removed with purge dups v1.2.5 (Guan *et al*. 2020). Mapping of Pacbio reads and Illumina WGS reads for the above steps were performed with minimap2 v2.17 and bwa-mem v0.7.17 respectively (Li 2018, 2013).

The trimmed HiC reads were aligned to the contig-level assembly with Juicer v1.6 (Durand *et al*. 2016). Scaffolding was performed with 3d-dna v180922 (Dudchenko *et al*. 2017). The initial scaffolding generated by 3d-dna was manually partitioned into chromosomes and misassembly corrected with Juicebox v1.11.08 (Robinson *et al*. 2018).

A k-mer spectrum, with *k* = 21 and a maximum counter value of 10^7^, was generated using KMC v3.1.1 (Kokot *et al*. 2017) and genome size was estimated from the spectrum using Genomescope v2.0 (Ranallo-Benavidez *et al*. 2020).

Gene completeness was evaluated using BUSCO v5.2.2 with the insecta odb10 dataset (n=1367) (Manni *et al*. 2021). Kmer QV was calculated using Merqury v1.3 (Rhie *et al*. 2020).

The mitochondrial genome was assembled and annotated using the Mitofinder pipeline v1.4 (Allio *et al*. 2020). Illumina WGS reads from SO_BI_364 were assembled with metaSPAdes v3.14.1 (Nurk *et al*. 2017) and tRNAs were annotated with MiTFi (Jühling *et al*. 2011).

### 2.4 Karyotype analysis

After scaffolding, chromosomes 11 and 13 displayed an intermediate HiC contact map pattern, suggesting a potential fusion of the chromosomes in one of the haplotypes.

In order to investigate this further we generated haplotype-specific HiC maps for chromosomes 11 and 13. First, we created a version of the assembly where chromosomes 11 and 13 were scaffolded together. WGS and HiC reads (from SO_BI_375) were mapped to this assembly with bwa-mem. Alignments were deduplicated with sambamba v0.6.6 (Tarasov *et al*. 2015). Heterozygous SNPs were called from the WGS alignments with freebayes v1.3.2-dirty (Garrison and Marth 2012), they were then normalised with bcftools v1.8 (Danecek *et al*. 2021), decomposed with vcfallelicprimitives (Garrison *et al*. 2021), and filtered for coverage (*>* 7 and *<* 56 reads) with bcftools. Filtered SNPs were phased using HAPCUT2 v1.3.3 with both the WGS and HiC alignments as input (Edge *et al*. 2017).

We developed a tool (chomper.py, see Data availability) which uses the phased SNPs from HAPCUT2 to partition aligned HiC reads by haplotype. For any read pair whose alignment encompasses at least one phased SNP we can ask whether the alleles in the read are associated with haplotype 1 or 2. If a read pair contains alleles exclusively associated with one haplotype then it is assigned to that haplotype-specific read set. If it instead contains alleles associated with both haplotypes then it is discarded. Haplotype-specific HiC read sets were then aligned back to the original assembly with Juicer and visualised with HiC_view.py (parameters `-b 250 -s 10`, see Data availability).

To identify the Z chromosome, one male (SO_BI_364) and one female (FR_BI_1497) individual were mapped to the assembly with bwa-mem and median, window-wise coverage was calculated using mosdepth v0.3.2 (Pedersen and Quinlan 2017).

### 2.5 Genome annotation

The Illumina RNA-seq reads were mapped to the assembly with HISAT2 v2.1.0 (Kim *et al*. 2019). The softmasked assembly and RNA-seq alignments were used for gene prediction with braker2.1.5 (Hoff *et al*. 2015, 2019; Li *et al*. 2009; Barnett *et al*. 2011; Lomsadze *et al*. 2014; Buchfink *et al*. 2015; Stanke *et al*. 2006, 2008). Gene annotation statistics were calculated with GenomeTools v1.6.1 (Gremme *et al*. 2013).

Transposable elements (TEs) were annotated using the Earl Grey TE annotation pipeline (https://github.com/TobyBaril/EarlGrey, Baril *et al*. 2021). Briefly, known repeats were masked with RepeatMasker v4.1.2 (Smit *et al*. 2015) using the Lepidoptera library from RepBase v23.08 and Dfam release 3.3 (Jurka *et al*. 2005; Hubley *et al*. 2015). Following this, a *de novo* repeat library was constructed using RepeatModeler2 v2.0.2 (Flynn *et al*. 2020) with RECON v1.08 and RepeatScout v1.0.6. Subsequently, Earl Grey generated maximum-length consensus sequences for the *de novo* sequences identified by RepeatModeler2 using an automated version of the ‘BLAST, Extract, Extend’ process, as previously described (Platt *et al*. 2016). The resulting *de novo* repeat library was combined with the RepBase and Dfam libraries used in the initial masking step to annotate repetitive elements using RepeatMasker. Full-length LTR elements were identified using LTR_Finder v1.07 with the LTR_Finder parallel wrapper (Xu and Wang 2007; Ou and Jiang 2019). Final TE annotations were defragmented and refined using a loose merge in RepeatCraft (`-loose`), followed by maintaining the longest of any overlapping annotations with MGkit v0.4.1 (`filter-gff -c length -a length`) (Wong and Simakov 2018; Rubino and Creevey 2014). Finally, all repeats *<*100bp in length were removed before final TE quantification to decrease spurious hits.

Following gene annotation, 5’ and 3’ flanking regions (’Gene Flanks’) were defined as those that were 20kb upstream and downstream of genes. TEs within these regions may overlap proximate or core promoter regions (Lenhard *et al*. 2012). We define regions as ’intergenic space’ if they are neither genic (start/stop codons, exons, and introns) nor Gene Flanks. Bedtools intersect v2.27.1 (Quinlan and Hall 2010) was used to determine overlap (`-wao`) between TEs and genomic features. Following this, quantification and plotting was performed in R, using the tidyverse package (Wickham *et al*. 2019; R Core Team 2021; RStudio Team 2020).

### 2.6 Estimating heterozygosity

To estimate heterozygosity, WGS reads were mapped to the assembly with bwa-mem and variants were called with freebayes. Callable sites, where coverage was *>* 7 and *<* twice the sample mean, were identified using mosdepth and fourfold-degenerate sites were identified using partition_cds.py (see Data availability). Variant calls were normalised with bcftools and decomposed using vcfallelicprimitives. Biallelic SNPs within callable fourfold-generate sites were counted using bedtools v2.30.0.

## 3 Results

### 3.1 Genome assembly

We sequenced and assembled the genome of a male *Brenthis ino* individual collected in Asturias, Spain (SO BI 364, Figures 1A and 1B). We generated 69.0-fold and 81.2-fold coverage of Pacbio CLR and Illumina WGS reads respectively. The initial assembly consisted of 119 contigs and had a total span of 411.8 Mb, which is consistent with a kmer-based estimate of haploid genome size of 414.0Mb (Figure S1). HiC reads (11.7-fold coverage) from a male specimen collected at the same locality (SO BI 375, Figures 1C and 1D) were used to scaffold the contigs into 14 chromosome-level sequences. These scaffolds range in size from 21.9 to 43.0 Mb and encompass 99.7% of the assembly. The contig and scaffold N50 of the assembly is 9.6 and 29.5 Mb respectively.

**Figure 1:**
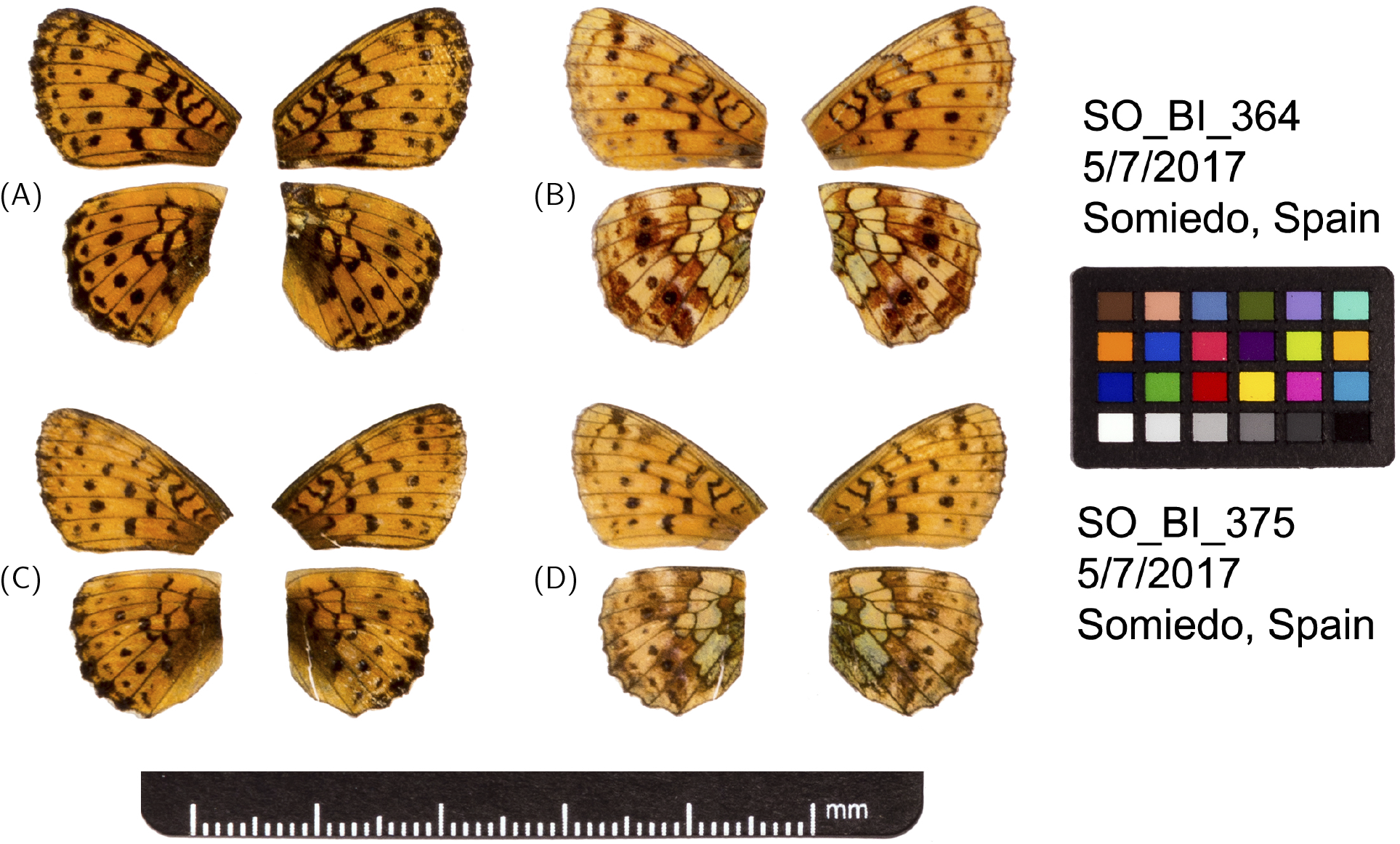
Fore and hind wings of the two *B. ino* individuals used to generate the genome sequence. (A) Dorsal and (B) ventral surface view of wings of specimen SO_BI_364, used to generate Pacbio and Illumina WGS reads. (C) Dorsal and (D) ventral surface view of wings of specimen SO_BI_375, used to generate HiC reads.

The BUSCO score of the assembly is 99.0% (S:98.6%, D:0.4%, F:0.3%, M:0.7%), suggesting the assembly is missing very few single-copy insect orthologues and has little duplication. The estimated mean Phred quality score of the consensus sequence is 39.85.

We assembled and annotated a circular mitochondrial genome of 15,180 bases with 13 protein coding genes, 22 tRNAs, and two rRNAs. The cytochrome oxidase subunit 1 (COI) nucleotide sequence has 99.85% identity (657/658 b) with a previously published COI sequence from a *B. ino* individual collected in Castilla y León, Spain (GenBank accession MN144802, Dapporto *et al*. 2019).

### 3.2 Evidence for a segregating neo-Z chromosome

While the HiC data support the scaffolding of 14 chromosome-level sequences (hereafter simply referred to as chromosomes), there is an excess of HiC contacts between chromosomes 11 and 13 (Figure 2A). This excess is not distributed evenly over the two chromosomes and is instead concentrated at one of the four possible junctions (Figure 2B), supporting the scaffolding of these two chromosomes in a specific orientation. However, while the number of HiC contacts between chromosomes 11 and 13 exceeds what we see between any other pair of chromosomes, it is below what we typically observe within chromosomes in this dataset (Figure S2), making it unclear whether chromosomes 11 and 13 are fused and should be scaffolded together. We tested whether the HiC contacts between chromosomes 11 and 13 are haplotype-specific, as this would result in half the number of contacts, and so could explain the reduced frequency (Figure S2). Haplotype-specific HiC maps (see Methods) confirm that HiC contacts between chromosomes 11 and 13 are almost entirely limited to one haplotype (Figures 2C and 2D) and the proportions of haplotype-specific reads (49.6% and 50.4% of partitioned reads support haplotypes 1 and 2 respectively) are consistent with these chromosomes being fused in one haplotype but not the other.

**Figure 2:**
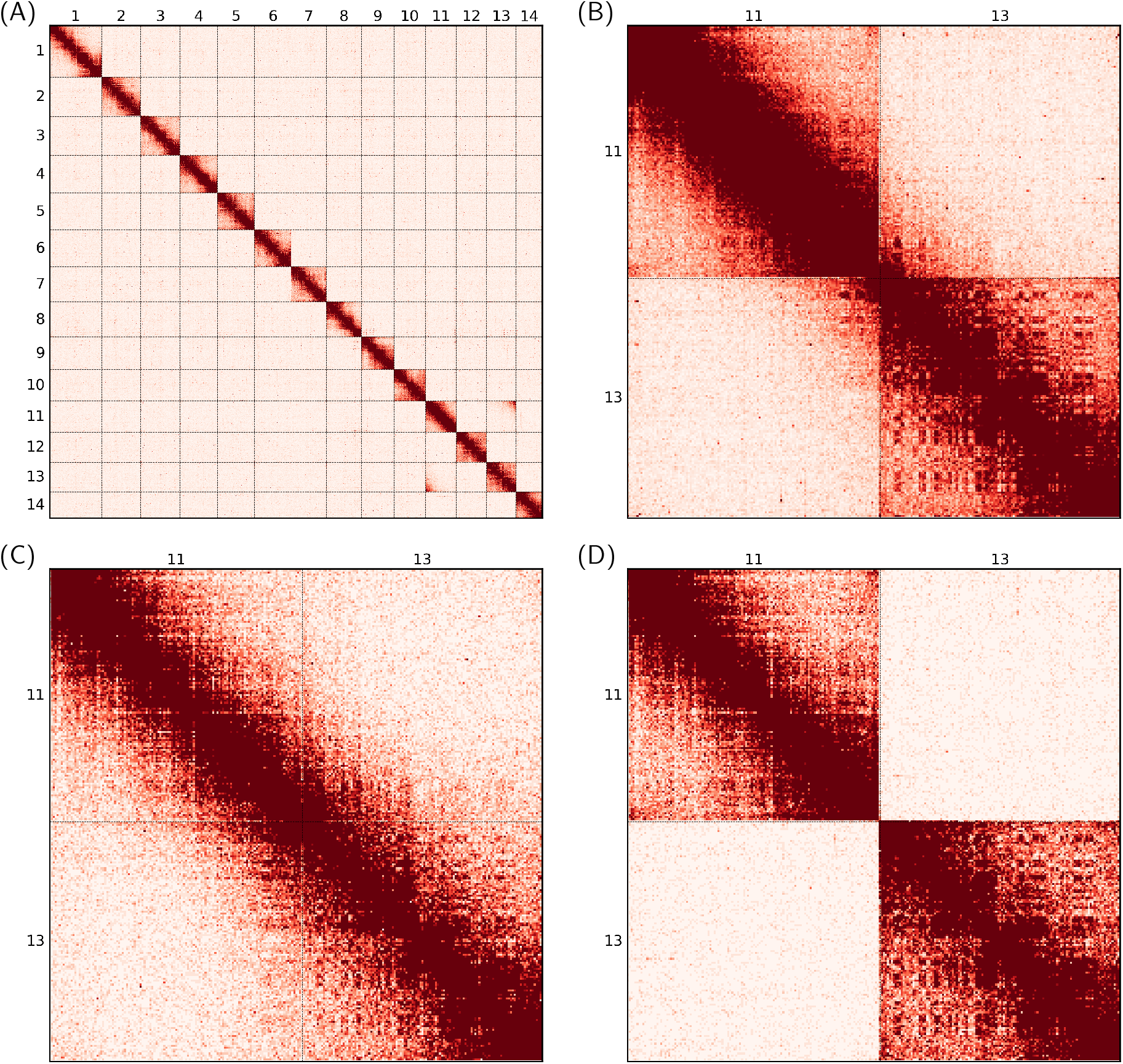
HiC contact heatmaps for the assembly of *B. ino*. (A) HiC contacts across all 14 chromosomes (HiC_view params: `-b 2500 -s` 25). (B) Contacts across chromosomes 11 and 13, with both chromosomes in the reverse orientation. (HiC_view params: `-b 250 -s` 30) (C) The same as in (B) but restricted to HiC reads containing alleles exclusively associated with haplotype 1 (HiC_view params: `-b 250 -s` 10). (D) The same as in (C) but associated with haplotype 2 rather than 1 (HiC_view params: `-b 250 -s` 10)

We identified chromosome 11 as the Z-chromosome in *B. ino*: the female individual (Figure S3) has half coverage for this chromosome, whereas the male used for assembly has full coverage (Figure S4). By contrast, chromosome 13 has full coverage in both males and females (Figure S4), consistent with the expectation for autosomal chromosomes (although see discussion). As one of these chromosomes is Z-linked, while the other has autosomal patterns of sex-specific coverage, we conclude that the individual from which we generated the HiC library must be heterozygous for a Z-autosome fusion, i.e. a neo-Z chromosome.

The Pacbio reads, which were generated from SO_BI_364 rather than SO_BI_375, do not span the gap between chromosomes 11 and 13. However, it is still possible that SO_BI_364 does possess a copy of the neo-Z chromosome, if the fusion point is within a region of the genome that is too repetitive to be assembled and so does not allow for successful chimeric alignment. It is therefore uncertain whether only SO_BI_375 possesses a copy of the neo-Z or if SO_BI_364 does as well.

### 3.3 Genome annotation

We annotated 16,844 protein coding genes. Given this annotation, we estimate that 33.5% of the genome assembly is intronic and 5.6% exonic. Chromosomes display some variation in gene density; chromosome 14, the shortest and most gene poor, is 32.8% genic whereas chromosome 11 (the Z) is 47.7% genic. Across the annotation, the median length of genes, introns, and exons is 4,084, 616 and 148 b, respectively.

Transposable elements (TEs) compromise 37.9% of the genome (Table S2, Figure 3A). Most TE activity appears to be relatively recent as a large proportion of repeats exhibit a low genetic distance from their respective consensus sequences (Figure 3B). The genome contains all major TE types (Table S2). Rolling circle elements, also known as helitrons, appear to have been the most successful progenitors within the genome, accounting for 17.8% of total genome span, and ∼ 47% of total TE content (Table S2). There is also evidence of very recent activity in LINEs and LTR elements, with a sharp increase in the number of identified elements with very low genetic distance to their consensus sequences (Figure 3B). The reasons for the recent bursts in LINEs and LTRs are unknown, although the likely recent age of these insertions is a sign of recent host colonisation, potentially via horizontal transposon transfer (HTT) from another host genome (Bourque *et al*. 2018; Oliveira *et al*. 2012; Ivancevic *et al*. 2018; Gilbert *et al*. 2010; Wallau *et al*. 2012).

**Figure 3:**
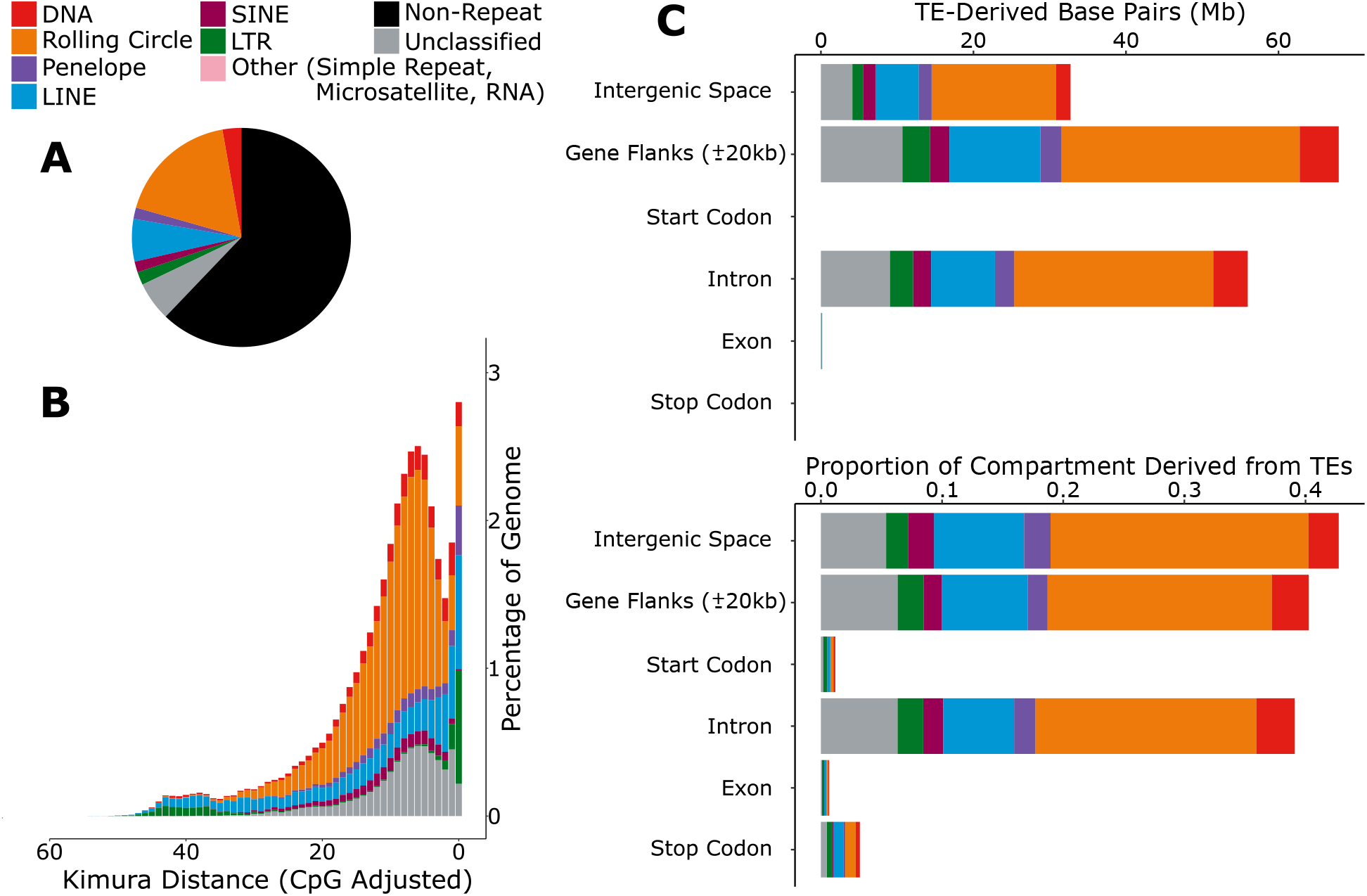
TEs within the genome assembly of *B. ino*. (A) The proportion of the assembly comprised of the main TE classifications, as represented by the colours in the key. (B) A repeat landscape showing the proportion of repeats with a given genomic distance to their consensus. More similar repeats are shown towards the right-hand side, indicating more recent activity. (C) The abundance of TEs in different partitions of the genome, shown in bases and as a proportion of the partition.

Considering all TE classifications, most TEs are found outside of genes (64.2% of total TE sequence: 20.9% in intergenic space, and 43.3% in gene flanking regions). Exons are largely devoid of TE sequence, with only 0.7% of exonic sequences consisting of TEs. This is to be expected given the likely detrimental effects of TE insertions in host exons (Bourque *et al*. 2018; Sultana *et al*. 2017). Meanwhile, intronic regions have amassed a higher proportion of TE sequences, resulting in 39.1% of total intronic sequence being derived from TEs (Figure 3C). The most abundant TEs in the genome, rolling circle elements, comprise 21.3% of intergenic space, ∼ 18% of gene flanking regions and intronic regions, and just 0.1% of exonic regions (Figure 3C).

## 4 Discussion

We have resolved the sequences of 14 *Brenthis ino* chromosomes: 13 autosomes and the Z sex-chromosome. The number of chromosomes in the assembly is higher than previously reported for *B. ino* in Europe (Federley 1938; Saitoh 1987, 1991) but equal to counts reported for this species in Japan (Maeki and Makino 1953; Saitoh *et al*. 1989). We note that previous karyotype data from Europe were all from Scandinavian samples, whereas the individuals contributing to the assembly were collected in Spain. It is therefore possible that the putative neo-Z, we describe here, exists at high frequency in Scandinavia, intermediate frequency in Spain, and low frequency in Japan, which could explain the contrasting karyotype observations for these populations.

We have interpreted the excess of HiC contacts between chromosomes 11 and 13, as well as the stark contrast in haplotype-specific HiC maps, as strong evidence for a segregating neo-Z chromosome. Lab contamination from a closely related - but karyotypically divergent - species is not a plausible alternative explanation given that the haplotype partitioned HiC reads are approximately equal in frequency (see Results). We can also rule out the possibility that we sampled an admixed individual, e.g. an F1 between *B. ino* and its sister species *B. daphne*, and that the neo-Z is fixed in one species but absent in the other. Both species are present in Northern Spain so sampling an F1 is possible, at least in principle. However, if SO_BI_375 were a recent hybrid, we would expect its heterozygosity to be considerably elevated compared to other *B. ino* individuals which is not the case: heterozygosity at autosomal fourfold degenerate sites for SO_BI_375, SO_BI_364, and FR_BI_1497, is 0.0109, 0.0106, and 0.0100, respectively, and in all cases is far lower than we would expect for an F1 between *B. ino* and *B. daphne* (∼ 0.025, Ebdon *et al*. 2021).

One way to further validate the existence of the neo-Z would be to test whether any females have half the normalised coverage over both chromosomes 11 and 13; which would be consistent with a single copy of the neo-Z (chromosomes 11 and 13 fused together), a W chromosome, but no additional copy of chromosome 13. However, if chromosome 13 is yet to evolve a dosage compensation mechanism, females carrying the neo-Z may only be viable with two copies of the autosomal sequence. Under this scenario, the female coverage seen in Figure S4 is consistent with both presence or absence of the neo-Z chromosome. Population level cytological or HiC data would be required to further validate the neo-Z and estimate its frequency.

While we have focused on karyotypic variation within a single *B. ino* individual, this assembly also provides an opportunity to test the causes and consequences of chromosome rearrangement accumulation more widely: at least 17 chromosome fusions must have happened on the lineage leading to *B. ino*, given an ancestral karyotype of 31 chromosomes (Ahola *et al*. 2014). Additionally, the *B. ino* assembly will enable population genomic studies in the genus *Brenthis*, expanding on previous reference-free population genomic analyses on this genus (Pazhenkova and Lukhtanov 2019; Ebdon *et al*. 2021). More generally, it adds to a growing number of high quality resources for comparative genomics in the Lepidoptera.

## 5 Data availability

Table S1 contains the metadata for the four individuals used for this project. The genome assembly, gene annotation, and raw sequence data can be found at the European Nucleotide Archive under project accession PRJEB49202. The scripts used for analysing HiC data (chomper.py and HiC_view.py) and the script used for calculating site degeneracy (partition_cds.py) can be found at the following github repository: https://github.com/A-J-F-Mackintosh/Mackintosh_et_al_2022_Bino. The mitochondrial genome sequence and the TE annotation can be found at the same repository.

## 6 Acknowledgments

We would like to thank Marian Thompson for preparing the Pacbio sequencing libraries, Karen Troup, Sarah White and Tony Miles for generating the Illumina libraries and Andres de la Filia and Katy MacDonald for help in the molecular lab. We also thank Maria Jesus Cañal Villanueva and Luis Valledor (Universidad de Orviedo) for help with fieldwork logistics. Collection permits for Somiedo were granted by the Gobierno del Principado de Asturias (014252) to KL. We thank Simon Martin for helpful discussion throughout the project and Sam Ebdon for taking the photos in Figure 1.

