## Supplementary material for "The genome sequence of the lesser marbled fritillary, *Brenthis ino*, and evidence for a segregating neo-Z chromosome"

### 9 Supplementary Materials

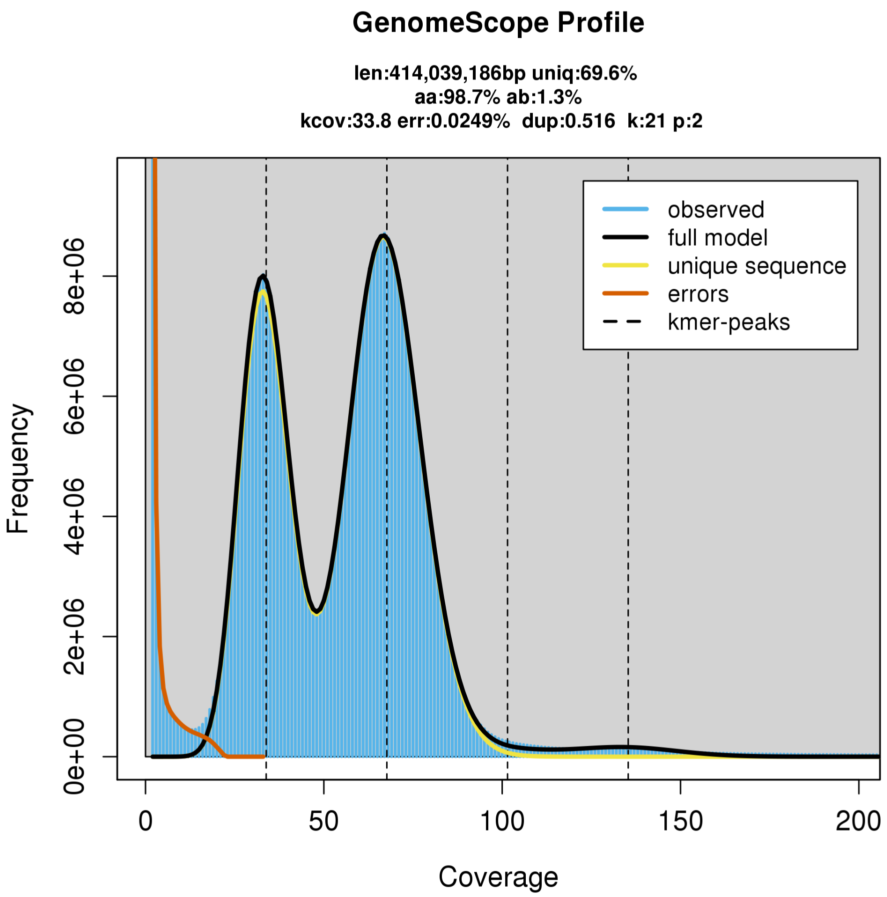


Figure S1: Kmer spectrum and Genomescope parameter estimates for SO_BI_364.

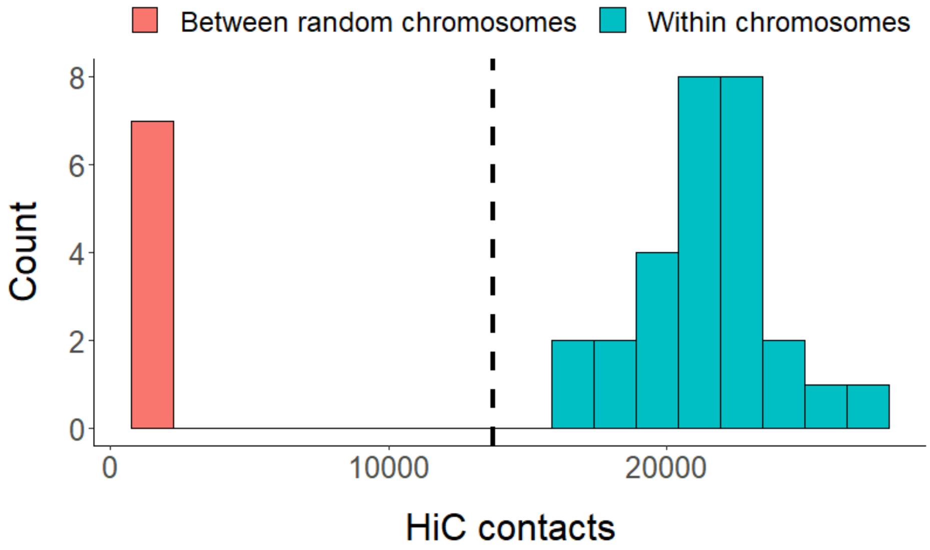


Figure S2: A histogram of HiC contact frequency within and between chromosomes. In red, the number of contacts spanning fusion points of randomly fused chromosome pairs, where either read is within 5Mb of the fusion point and each chromosome has been randomly fused once. In blue, the number of contacts spanning arbitrary points within chromosomes, where reads are again within 5Mb and two independent points are sampled per chromosome. The dashed line represents the number of contacts spanning the putative neo-Z fusion between chromosomes 11 and 13. The frequency of contacts supporting the neo-Z is consistent with heterozygosity.

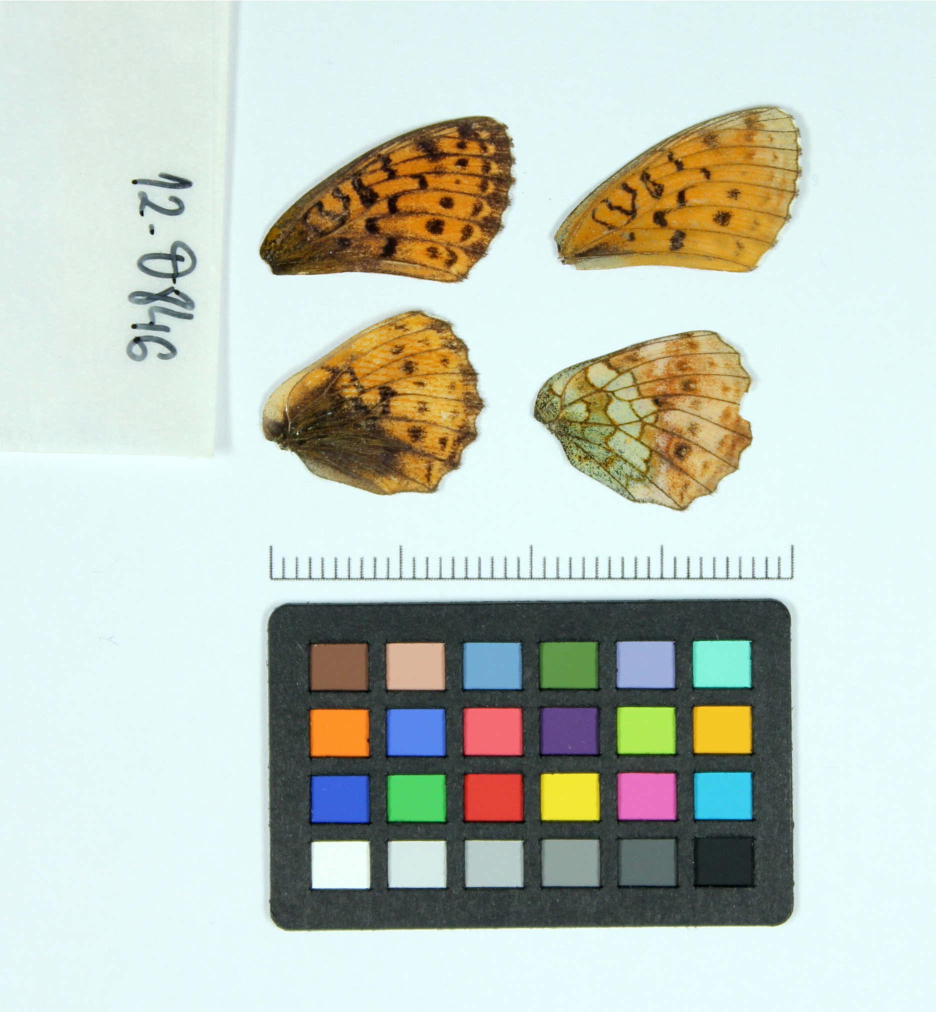

Figure S3: Wings of the female specimen FR_BI_1497. Top-left: dorsal forewing. Top-right: ventral forewing. Bottom-left: dorsal hindwing. Bottom-right: ventral hindwing

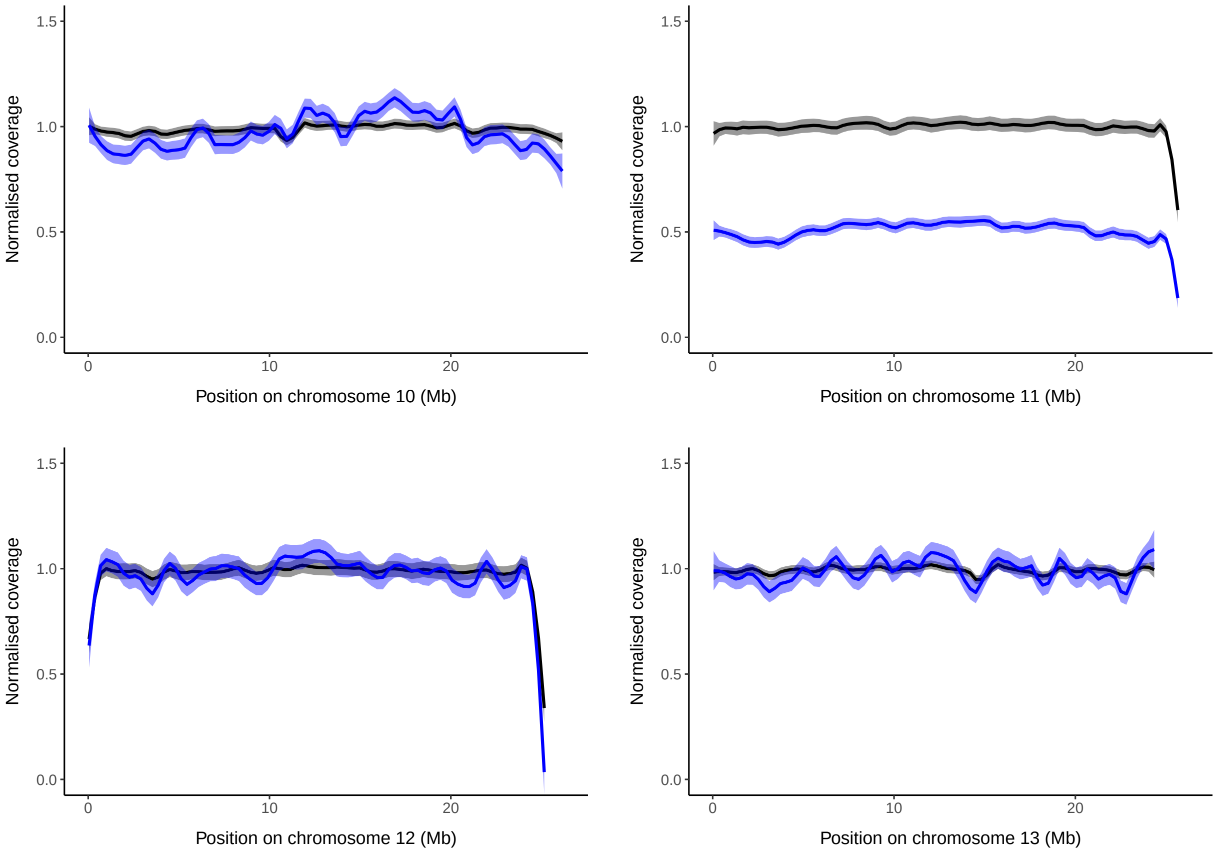


Figure S4: Normalised coverage on four chromosomes. The black line represents normalised coverage of WGS reads from the male used for genome assembly (SO_BI_364), while the blue line is normalised coverage of WGS reads from a female individual (FR_BI_1497). On chromosome 11 the male has full coverage whereas the female has half coverage, consistent with expectations for Z-linked chromosomes in Lepidoptera. On the other three chromosomes, as well as the ten not shown, both individuals have full normalised coverage.

Table S1: Sampling locations and other metadata for *B. ino* samples used in this study.

| Sample | Date | Sex | Locality | Region | Country | Lat | Long | Altitude | Collector |
|---|---|---|---|---|---|---|---|---|---|
| SO_BI_364 | 05/07/2017 | Male | Somiedo, Braña de Mumian | Asturias | Spain | 43.068 | -6.24 | 1420 | KL |
| SO_BI_375 | 05/07/2017 | Male | Somiedo, Braña de Mumian | Asturias | Spain | 43.068 | -6.24 | 1420 | KL |
| SO_BI_376 | 05/07/2017 | n/a | Somiedo, Braña de Mumian | Asturias | Spain | 43.068 | -6.24 | 1420 | KL |
| FR_BI_1497 (RV-coll12O846) | 11/08/2012 | Female | Larche (Les Marmottes) | Alpes-de-Haute-Provence | France | 44.446 | 6.851 | 1680 | VD & Raluca Vodă |

Table S2: Annotated transposable elements

| Repeat class | No. elements | Total length (Mb) | Percentage of genome (%) | No. distinct classifications |
|---|---|---|---|---|
| **Retroelement** | 116256 | 47.36 | 11.49 | 930 |
| SINE | 26242 | 6.53 | 1.58 | 23 |
| LINE | 59472 | 25.92 | 6.29 | 611 |
| Penelope | 25145 | 6.89 | 1.67 | 34 |
| LTR element | 5397 | 8.02 | 1.95 | 262 |
| **DNA transposon** | 36793 | 11.48 | 2.79 | 598 |
| **Rolling Circle** | 258724 | 73.34 | 17.81 | 307 |
| **Unclassified** | 74790 | 23.74 | 5.76 | 339 |
| **Other** | 21 | 0.00 | 0.00 | 3 |
| **Total** | 486584 | 155.92 | 37.85 | 2177 |
